# Inferotemporal cortex multiplexes behaviorally-relevant target match signals and visual representations in a manner that minimizes their interference

**DOI:** 10.1101/152181

**Authors:** Noam Roth, Nicole C. Rust

**Affiliations:** Department of Psychology, University of Pennsylvania, Philadelphia, PA 19104

## Abstract

Finding a sought visual target object requires combining visual information about a scene with a remembered representation of the target to create a “target match” signal that indicates when a target is in view. Target match signals have been reported to exist within high-level visual brain areas including inferotemporal cortex (IT), where they are mixed with representations of image and object identity. However, these signals are not well understood, particularly in the context of the real-world challenge that the objects we search for typically appear at different positions, sizes, and within different background contexts. To investigate these signals, we recorded neural responses in IT as two rhesus monkeys performed a delayed-match-to-sample object search task in which target objects could appear at a variety of identity-preserving transformations. Consistent with the existence of behaviorally-relevant target match signals in IT, we found that IT contained a linearly separable target match representation that reflected behavioral confusions on trials in which the monkeys made errors. Additionally, target match signals were highly distributed across the IT population, and while a small fraction of units reflected target match signals as target match suppression, most units reflected target match signals as target match enhancement. Finally, we found that the potentially detrimental impact of target match signals on visual representations was mitigated by target match modulation that was approximately (albeit imperfectly) multiplicative. Together, these results support the existence of a robust, behaviorally-relevant target match representation in IT that is configured to minimally interfere with IT visual representations.

## Introduction

Finding a sought visual target object requires combining incoming visual information about the identities of the objects in view with a remembered representation of a sought target object to create a “target match” signal that indicates when a target has been found. During visual target search, target match signals have been reported to emerge in the brain as early as visual areas V4 (Bichot et al., 2005; Chelazzi et al., 2001; Haenny et al., 1988; Kosai et al., 2014; Maunsell et al., 1991) and IT (Chelazzi et al., 1998; Chelazzi et al., 1993; Eskandar et al., 1992; Gibson and Maunsell, 1997; Leuschow et al., 1994; Mruczek and Sheinberg, 2007; Pagan et al., 2013; Woloszyn and Sheinberg, 2009). However, we understand very little about the nature of target match signals, their behavioral relevance, and how these signals are mixed with visual representations.

The nature of the target match signal has been investigated most extensively with traditional versions of the delayed-match-to-sample (DMS) paradigm, which involves the presentation of a cue image indicating a target’s identity, followed by the presentation of a random number of distractors and then a target match (e.g. Haenny et al., 1988; Miller and Desimone, 1994; Pagan et al., 2013). During classic DMS tasks in which the cue is presented at the beginning of each trial (and the match is thus a repeat later on), IT has been reported to reflect target match information with approximately equal numbers of neurons preferring target matches versus those preferring distractors (i.e. “target match enhancement” and “target match suppression”, respectively; Miller and Desimone, 1994; Pagan et al., 2013). Upon observing that target match suppression also follows from the repetition of distractor images within a trial, and thus cannot account for a signal that corresponds to a “target match” behavioral report, some have speculated that target match enhancement alone reflects the signal used to make behavioral judgments about whether a target match is present (Miller and Desimone, 1994). Others have proposed that the responses of both target match enhanced and suppressed subpopulations are incorporated to make behavioral judgments, particularly when a task requires disambiguating changes in firing rate due to the presence of a target match from other factors that impact overall firing rate, such as stimulus contrast (Engel and Wang, 2011). Notably, no study to date has produced compelling evidence that either IT target match enhancement or suppression accounts for (or correlates with) behavioral reports (e.g. on error trials).

Another limitation of the traditional DMS paradigm is that the cue image tends to be an exact copy of the target match, whereas real-world object search involves searching for an object that can appear at different positions, sizes and background contexts. One DMS study examined IT neural responses during this type of object variation and reported the existence of target match signals under these conditions (Leuschow et al., 1994). However, we still do not understand how IT target match signals are intermingled with IT invariant object representations of the currently-viewed scene. One intriguing proposal (Fig 1) suggests how visual and target match signals might be multiplexed to minimize the interference between them. That is, insofar as visual representations of different images are reflected as distinct patterns of spikes across the IT population (reviewed by DiCarlo et al., 2012), this translates into a population representation in which visual information is reflected by the population vector angle (Fig 1, ‘Visual modulation’). If the introduction of target match modulation also changes population vector angles in IT, this could result in perceptual confusions about the visual scene. However, if target match modulation amounts to multiplicative rescaling of population response vector lengths, this would minimize interference when superimposing visual memories and target match representations within the same network (Fig 1, ‘Target match modulation’). The degree to which the target match signal acts in this way remains unknown.

**Figure 1.**
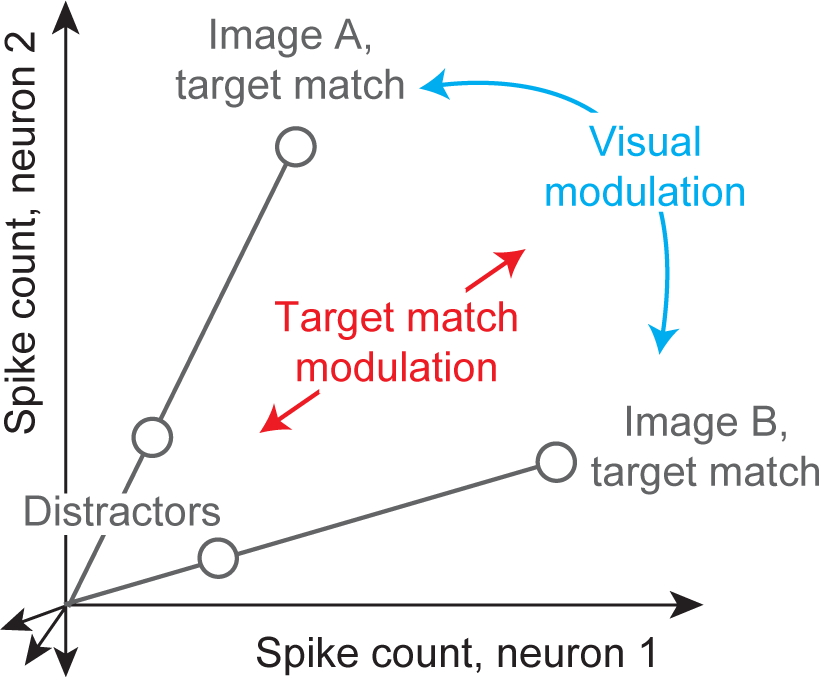
Multiplexing visual and target match representations. Shown are the hypothetical population responses to two images, each viewed (at different times) as target matches versus as distractors, plotted as the spike count response of neuron 1 versus neuron 2. In this scenario, visual information (e.g. image or object identity) is reflected by the population response pattern, or equivalently, the angle that each population response vector points. In contrast, target match information is reflected by changes in population vector length (e.g. multiplicative rescaling). Because target match information does not impact vector angle in this hypothetical scenario, superimposing target match information in this way would mitigate the impact of intermingling target match signals within underlying perceptual representations.

To investigate the nature of the IT target match signal, its behavioral relevance, and how it intermingles with IT visual representations, we recorded neural signals in IT as monkeys performed a modified delayed-match-to-sample task in which they were rewarded for indicating when a target object appeared across changes in the objects’ position, size and background context.

## Results

### The invariant delayed-match-to-sample task (IDMS)

To investigate the target match signal, we trained two monkeys to perform an “invariant delayed-match-to-sample” (IDMS) task that required them to report when target objects appeared across variation in the objects’ positions, sizes and background contexts. In this task, the target object was held fixed for short blocks of trials (~3 minutes on average) and each block began with a cue trial indicating the target for that block (Fig 2a, “Cue trial”). Subsequent test trials always began with the presentation of a distractor and on most trials this was followed by additional distractors and then an image containing the target match (Fig 2a, “Test trial”). The monkeys’ task required them to fixate during the presentation of distractors and make a saccade to a response dot on the screen following target match onset to receive a reward. In cases where the target match was presented for 400 ms and the monkey had still not broken fixation, a distractor stimulus was immediately presented. To minimize the predictability of the match appearing as a trial progressed, on a small subset of the trials the match did not appear and the monkey was rewarded for maintaining fixation. Our experimental design differs from other classic DMS tasks (e.g. Miller and Desimone, 1994; Pagan et al., 2013) in that it does not incorporate a cue at the beginning of each test trial, to better mimic real-world object search conditions in which target matches are not repeats of the same image presented shortly before.

**Figure 2.**
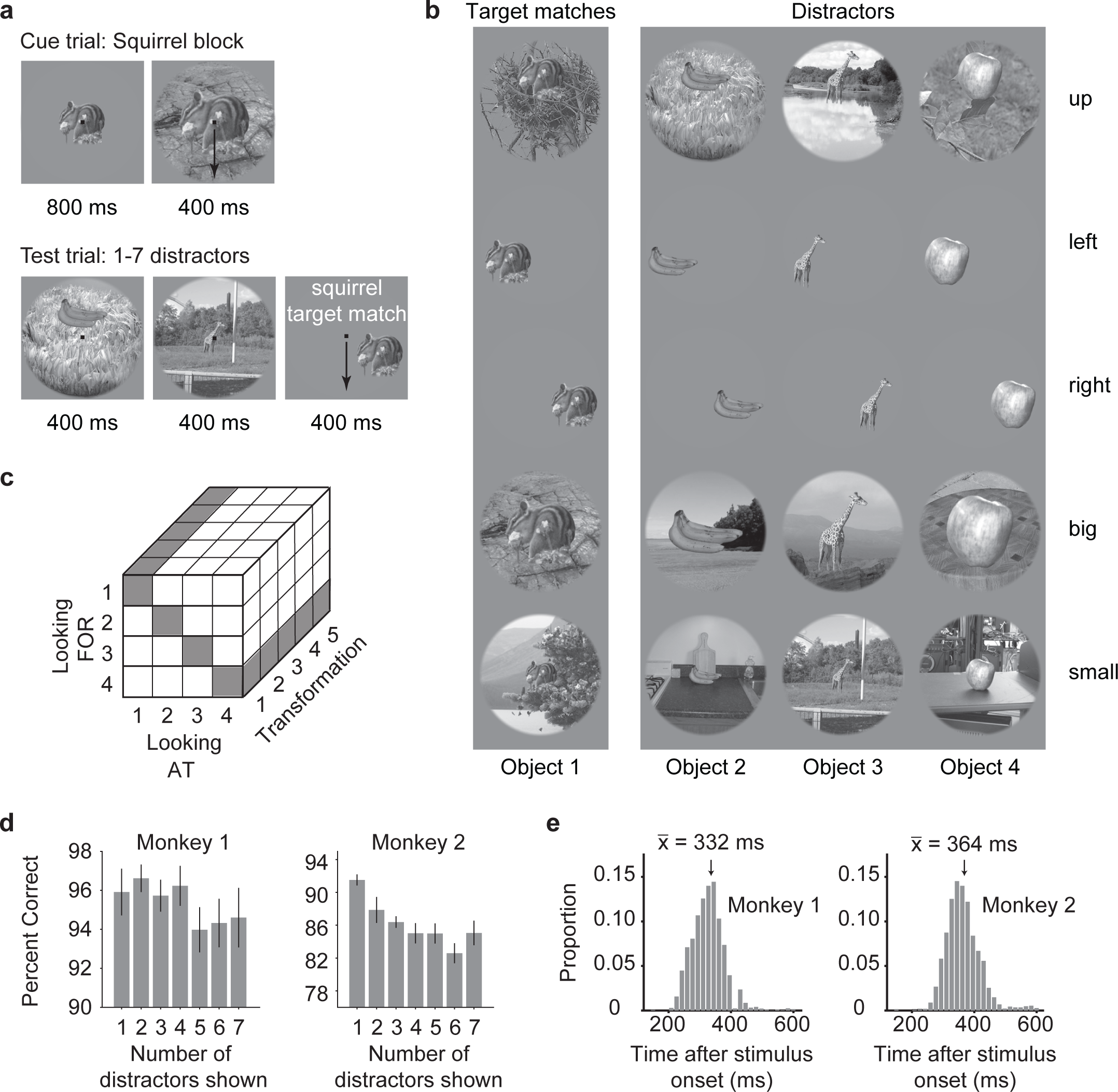
The invariant delayed-match-to-sample task. **a)** Each block began with a cue trial indicating the target object for that block. On subsequent trials, no cue was presented and monkeys were required to maintain fixation throughout the presentation of distractors and make a saccade to a response dot following the onset of the target match to receive a reward. **b)** The experiment included 4 objects presented at each of 5 identity-preserving transformations (“up”, “left”, “right”, “big”, “small”), for 20 images in total. In any given block, 5 of the images were presented as target matches and 15 were distractors. **c)** The complete experimental design included looking “at” each of 4 objects, each presented at 5 identity-preserving transformations (for 20 images in total), viewed in the context of looking “for” each object as a target. In this design, target matches (highlighted in gray) fall along the diagonal of each “looking at” / “looking for” transformation slice. **d)** Percent correct for each monkey, calculated based on both misses and false alarms (but disregarding fixation breaks). Mean percent correct is plotted as a function of the position of the target match in the trial. Error bars (SEM) reflect variation across the 20 experimental sessions. e) Histograms of reaction times during correct trials (ms after stimulus onset) during the IDMS task for each monkey, with means indicated by arrows and labeled.

Our experiment included a fixed set of 20 images, including 4 objects presented at each of 5 transformations (Fig 2b). Our goal in selecting these specific images was to make the task of classifying object identity challenging for the IT population and these specific transformations were built on findings from our previous work (Rust and DiCarlo, 2010). In any given block (e.g. a squirrel target block), a subset of 5 of the images would be considered target matches and the remaining 15 would be distractors (Fig 2b). Our full experimental design amounted to 20 images (4 objects presented at 5 identity-preserving transformations), all viewed in the context of each of the 4 objects as a target, resulting in 80 experimental conditions (Fig 2c). In this design, “target matches” fall along the diagonals of each looking at / looking for matrix slice (where “slice” refers to a fixed transformation; Fig 2c, gray). For each condition, we collected at least 20 repeats on correct trials. Monkeys generally performed well on this task (Fig 2d; mean percent correct monkey 1 = 96%; monkey 2 = 87%). Their mean reaction times (computed as the time their eyes left the fixation window relative to the target match stimulus onset) were 332 ms and 364 ms (Fig 2e).

As two monkeys performed this task, we recorded neural activity in IT using 24-channel probes. We performed two types of analyses on these data. The first type of analysis was performed on the data recorded simultaneously across units within a single recording session (n=20 sessions, including 10 sessions from each monkey). The second type of analysis was performed on data that was concatenated across different sessions to create a pseudopopulation after screening for units based on their stability, isolation, and task modulation (see Methods; n=204 units in total, including 108 units from monkey 1 and 96 units from monkey 2; S1 Dataset). For all but four of our analyses (Fig 4b, 4d, 8, 9), we counted spikes in a window that started 80 ms following stimulus onset (to allow stimulus-evoked responses time to reach IT) and ended at 250 ms, which was always before the monkeys’ reaction times on these trials. For all but two of our analyses (Fig 6, 7d), the data are extracted from trials with correct responses.

### Target match signals were reflected in IT during the IDMS task

Distributions of stimulus-evoked firing rates for the 204 units recorded in our experiment are shown in Figure 3. As is typical of IT and other high-level brain areas, we encountered a heterogeneous diversity of units with regard to their tuning to different aspects of the IDMS task. Figure 4a depicts the responses of four example units, plotted as five slices through our experimental design matrix (Fig 2c), where each slice corresponds to viewing each of the four objects at a fixed transformation (‘Looking AT’) in the context of searching for each of the four objects as a target (‘Looking FOR’). Different types of task modulation produce distinct structure in these response matrices. Visual modulation translates to vertical structure, (e.g. looking at the same image while looking for different things) whereas target modulation translates to horizontal structure (e.g. looking for the same object while looking at different things). In contrast, target match modulation is reflected as a differential response to the same images presented as target matches (which fall along the diagonal of each slice) versus distractors (which fall off the diagonal), and thus manifests as diagonal structure in each slice.

**Figure 3.**
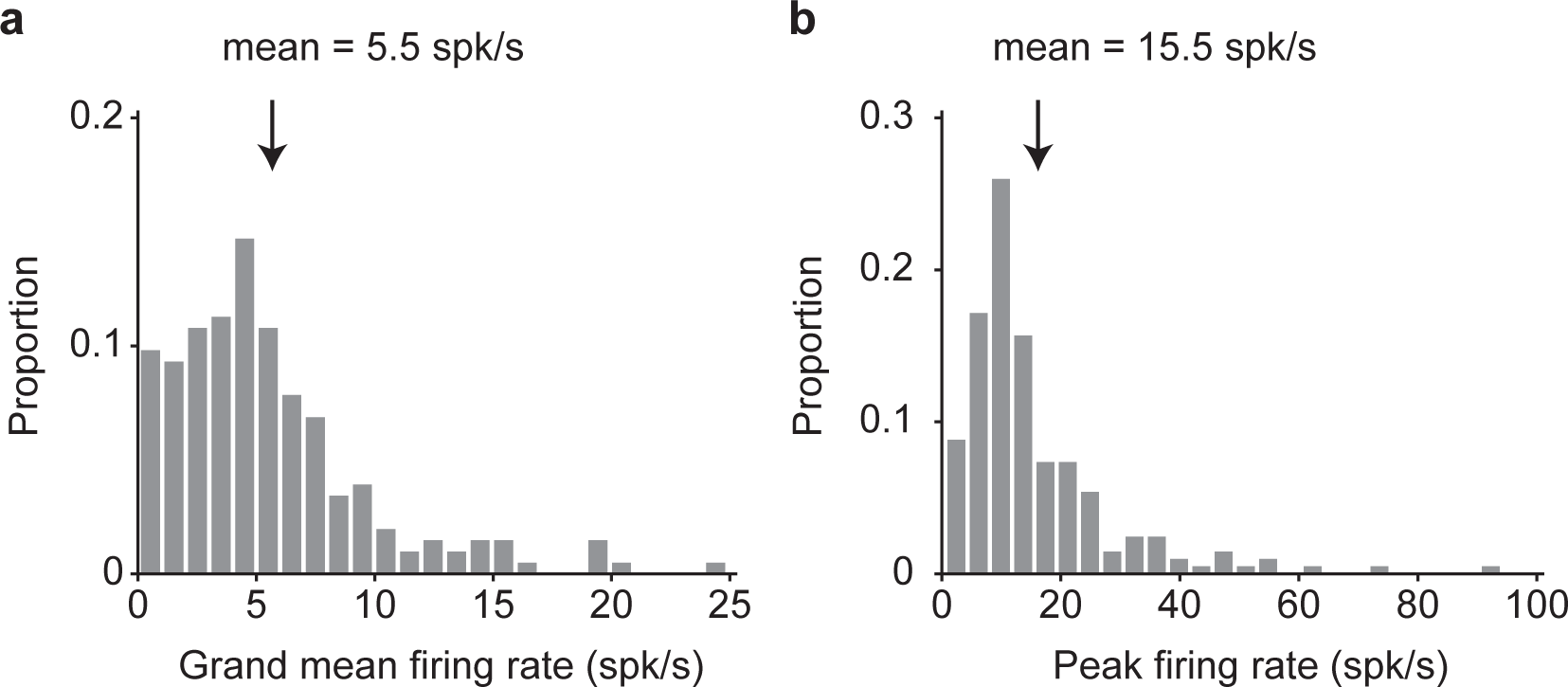
Firing rate distributions. The firing rate response to each stimulus was calculated as the mean across 20 trials in a window 80 - 250 ms following stimulus onset. **a)** Grand mean firing rate across all 80 conditions. **b)** Maximum firing rates across the 80 conditions. Arrows indicate the means (n=204 units).

**Figure 4.**
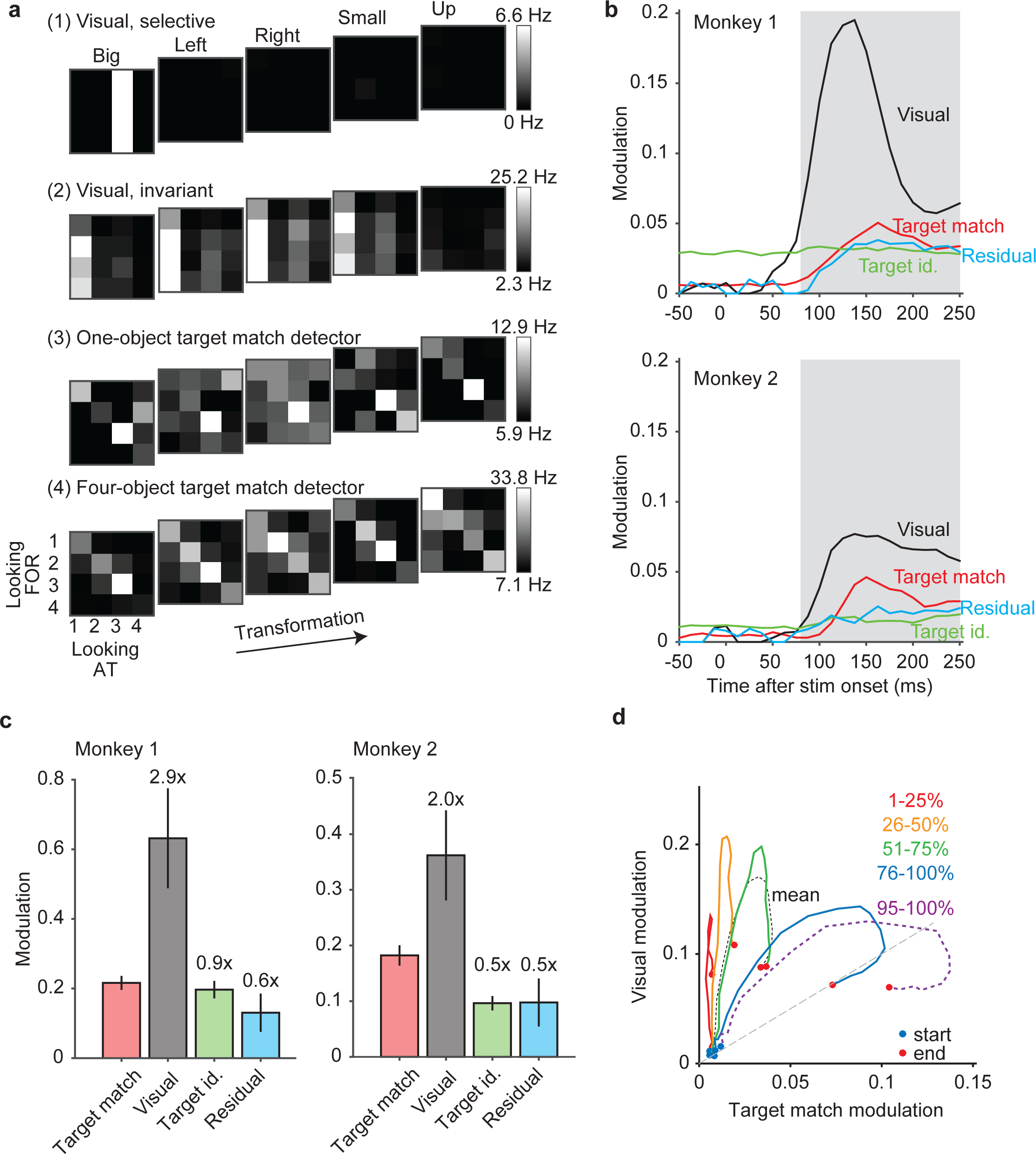
Quantifying modulation in IT during the IDMS task. **a)** The response matrices corresponding to four example IT units, plotted as the average response to five slices through the experimental design, where each slice (a 4×4 matrix) corresponds to viewing each of four objects (‘Looking AT’) in the context of each of four objects as a target (‘Looking FOR’), at one transformation (‘Big’, ‘Left’, ‘Right’, ‘Small’, ‘Up’). To compute these responses, spikes were counted in a window 80 −250 ms following stimulus onset, averaged across 20 repeated trials, and rescaled from the minimum (black) to maximum (white) response across all 80 conditions. **b)** Firing rate modulations were parsed into constituent types, where modulation was quantified in units of standard deviation around each unit’s grand mean spike count (see Results). The evolution of average modulation magnitudes (across all the units for each animal; monkey 1: n = 108, monkey 2: n = 96), shown as a function of time relative to stimulus onset. The shaded area indicates the spike count window used for subsequent analyses. **c)** Average modulation magnitudes computed using the spike count window depicted in panel b. **d)** The average temporal evolution of visual modulation plotted against the average temporal evolution of target match modulation for groups of units organized into quantiles. Units with either target match or visual modulation (n=203 of 204 units) were sorted by their ratios of target match over visual modulation, computed in a window 80-250 ms following stimulus onset. The temporal evolution of the mean across the population (black dotted line) is plotted among the temporal evolution of each 25% quartile of the data, as well as the 95-100% quantile (labeled). Start times of each trajectory (0 ms after stimulus onset) are indicated by a blue dot whereas end times of each trajectory (250 ms after stimulus onset) are indicated by a red dot.

The first example unit (Fig 4a, ‘visual, selective’) only responded to one image (object 3 presented in the “big” transformation) and was unaffected by target identity. In contrast, the second example unit (‘Fig 4a, ‘visual, invariant’) responded fairly exclusively to one object, but did so across four of the five transformations (all but “up”). This unit also had modest target match modulation, reflected as a larger response to its preferred object (object 1) when it was a distractor (i.e. when searching for targets 2-4) as compared to when it was a target (i.e. when searching for target 1). In other words, this unit exhibited target match suppression. The third example unit (“Fig 4a, ‘one-object target match detector’) consistently responded with a high firing rate to object 3 presented as a target match across all transformations, but not to other objects presented as target matches. This unit thus exhibited a form of target match enhancement that was selective for object identity. The fourth example unit (“Fig 4a, ‘four-object target match detector’) responded in a compelling way with a higher firing rate response to nearly any image (any object at any transformation) presented as a target match as compared to as a distractor, or equivalently target match enhancement that was invariant to object identity. Given that the IDMS task requires an eye movement in response to images presented as target matches and fixation to the same images presented as distractors, this unit reflects something akin to the solution to the monkeys’ task.

To quantify the amounts of these different types of modulations across the IT population, we applied a procedure that quantified different types of modulation in terms of the number of standard deviations around each unit’s grand mean spike count (Pagan and Rust, 2014b). Our procedure includes a bias-correction to ensure that modulations are not over-estimated by trial variability and it is similar to a multi-way ANOVA, with important extensions (see Methods). Modulation magnitudes were computed for the types described above, including visual, target identity, and target match modulation, as well as “residual” modulations that are reflected as nonlinear interactions between the visual stimulus and the target identity that are not captured by target match modulation (e.g. specific distractor conditions). Notably, this analysis defines target match modulation as a differential response to the same images presented as target matches versus distractors, or equivalently, diagonal structure in the transformation slices presented in Fig 4a. Consequently, units both like the “one-object target match detector” as well as the “four-object target match detector” reflect target match modulation, as both units have a diagonal component to their responses. What differentiates these two units is that the “one-object target match detector” also reflects selectivity for image and target identity, reflected in this analysis as a mixture of target match, visual, and target identity modulation.

Figure 4b illustrates these modulations computed in a sliding window relative to stimulus onset and averaged across all units recorded in each monkey. As expected from a visual brain area, we found that visual modulation was robust and delayed relative to stimulus onset (Fig 4b, black). Visual modulation was considerably larger in monkey 1 as compared to monkey 2. Target match modulation (Fig 4b, red) was also (as expected) delayed relative to stimulus onset and was smaller than visual modulation, but it was well above the level expected by noise (i.e. zero) and was similar in magnitude in both animals. In contrast, a robust signal reflecting information about the target identity (Fig 4b, green) appeared before stimulus onset in monkey 1 and was weaker but also present in monkey 2, consistent with a persistent working memory representation. Note that because the IDMS task was run in blocks with a fixed target, target identity information was in fact present before the onset of each stimulus. Lastly, we found that residual modulation was also present but was smaller than target match modulation in both animals (Fig 4b, cyan). In sum, for a brief period following stimulus onset, visual and target signals were present, but target match signals were not. After a short delay, target match signals appeared as well. When quantified in a window positioned 80 to 250 ms following stimulus onset and computed relative to the size of the target match signal (Fig 4c), visual modulation was considerably larger than target match modulation (monkey 1: 2.9x, monkey 2: 2.0x; Fig 4c, gray), whereas the other types of modulations were smaller than target match modulation (target modulation, monkey 1: 0.9x, monkey 2: 0.5x, Fig 4c green; residual modulation, monkey 1: 0.6x, monkey 2: 0.9x Fig 4c, cyan).

To what degree do these population average traces (Fig 4b) reflect the evolution of visual and target match signals in the same units as opposed to different units? To address this question, we ranked units by their ratios of target match and visual modulation, and grouped them into quantiles of neighboring ranks. Fig 4d shows a plot of the temporal evolution of visual modulation plotted against the evolution of target match modulation for each 25% quartile. The lowest-ranked quartile (Fig 4d, red) largely traversed and then returned along the y-axis, consistent with units that were nearly completely visually modulated. Of interest was whether quartiles with higher ratios of target match modulation would traverse the x-axis in an analogous fashion (reflecting pure target match modulation) or whether these units would begin as visually modulated and become target match modulated at later times. The trajectories for all three higher quartiles (Fig 4d, orange, green, blue) reflected the latter scenario, as they all began with a visually dominated component positioned above the unity line (Fig 4d, gray dashed). Later, the trajectories become more horizontal, indicative of the emergence of target match modulation. Similarly, the trajectory confined to just the top 5% (n=8) units (Fig 4d, purple dashed) began with a visually dominated component that later evolved into strong target match modulation. These results suggest that the evolution of visual to target match modulation is not happening within distinct subpopulations, but rather is reflected within individual units.

To summarize, the results presented thus far verify the existence of a target match signal in IT that is on average ~40% of the size of the visual modulation. Additionally, while the arrival of target match modulation was delayed relative to the arrival of visual modulation, both types of modulation tend to be reflected in the same units (as opposed to distinct subpopulations).

### IT target match information was reflected as a highly distributed, linearly separable representation

The IDMS task required monkeys to determine whether each condition (an image viewed in the context of a particular target block) was a target match or a distractor. This task ultimately maps all the target match conditions onto one behavioral response (a saccade) and all the distractor conditions onto another (maintain fixation), and as such, this task can be envisioned as a twoway classification across changes in other parameters, including changes in target and image identity (Fig 5a). One question of interest is the degree to which the target match versus distractor classification can be made with a linear decision boundary (or equivalently a linear decoder) applied to the IT neural data, as opposed to requiring a nonlinear decoding scheme. In a previous study, we assessed the format of IT target match information in the context of the classic DMS task design (Pagan et al., 2016; Pagan et al., 2013) and found that while a large component was linear, a considerable nonlinear (quadratic) component existed as well.

To quantify the amount and format of target match information within IT, we began by quantifying cross-validated performance for a two-way target match versus distractor classification with a weighted linear population decoder (a Fisher Linear Discriminant, FLD). Linear decoder performance began near chance and grew as a function of population size, consistent with a robust IT target match representation (Fig 5b, white). To determine the degree to which a component of IT target match information was present in a nonlinear format that could not be accessed by a linear decoder, we measured the performance of a maximum likelihood decoder designed to extract target match information regardless of its format (combined linear and nonlinear, Pagan et al., 2016; Pagan et al., 2013, see Methods). Performance of this nonlinear decoder (Fig 5b, gray) was slightly higher than the linear decoder for the pooled data (p = 0.022), and was not consistently higher in both animals (monkey 1 p = 0.081; monkey 2 p = 0.647). These results suggest that under the conditions of our measurements (e.g. the population sizes we recorded and the specific images used), IT target match information is reflected almost exclusively in a linearly separable format during the IDMS task. These results are at apparent odds with our previous reports of how IT target match information is reflected during a classic DMS task (see Discussion).

**Figure 5.**
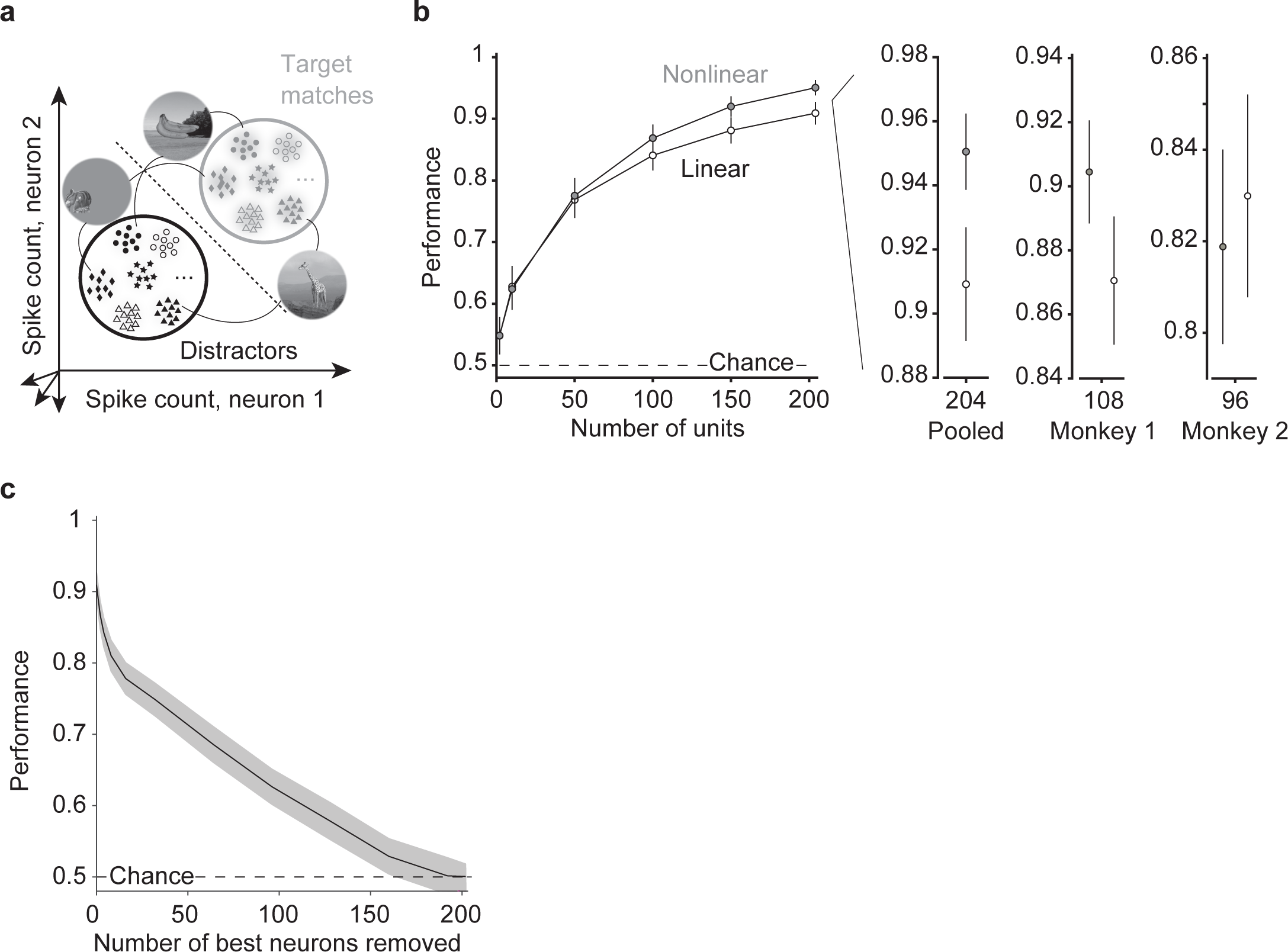
IT target match information is reflected via weighted linear scheme. **a)** The target search task can be envisioned as a two-way classification of the same images presented as target matches versus as distractors. Shown are cartoon depictions where each point depicts a hypothetical response of a population of two neurons on a single trial, and clusters of points depict the dispersion of responses across repeated trials for the same condition. Included are responses to the same images presented as target matches and as distractors. Here only 6 images are depicted but 20 images were used in the actual analysis. The dotted line depicts a hypothetical linear decision boundary. **b)** Linear (FLD) and nonlinear (maximum likelihood) decoder performance as a function of population size for a pseudopopulation of 204 units combined across both animals, as well as for the data recorded in each monkey individually (monkey 1: n = 108 units; monkey 2: n = 96 units.) Error bars (SEM) reflect the variability that can be attributed to the random selection of units (for populations smaller than the full dataset) and the random assignment of training and testing trials in cross-validation. **c)** Linear (FLD) decoder performance as a function of the number of top-ranked units removed. Shaded error (SEM) reflects the variability that can be attributed to the random assignment of training and testing trials in cross-validation.

Next, we wanted to better understand how target match information was distributed across the IT population. We thus performed an analysis targeted at the impact of excluding the N “best” target match units for different values of N, with the rationale that if it were the case that the majority of target match information was carried by a small subpopulation of units, performance should fall quickly when those units are excluded. For this analysis, we considered the magnitude but not the sign of the target match modulation (whereas we address questions related to parsing target match modulation by sign, or equivalently target match enhancement versus suppression, below in Figure 7). To perform this analysis, we excluded the top-ranked IT units via a cross-validated procedure (i.e. based on the training data; see Methods). Consistent with a few units that carry target match signals that are considerably stronger than the rest of the population, we found that the slope of the performance drop following the exclusion of the best units was steepest for the top 8% (n=16) ranked units, and that these units accounted for ~25% of total population performance (Fig 5c). However, it was also the case that population performance continued to decline steadily as additional units were excluded, and consequently, population performance could not be attributed to a small fraction of top-ranked units alone (Fig 5c). For example, a 50% decrement in performance required removing 27% (n=55/204) of the best-ranked IT population, and mean +/- SEM performance remained above chance up to the elimination of 78% (n=160/204) of top-ranked units. These results are consistent with target match signals that are strongly reflected in a few units (such as Fig 4a example unit 4), and are more modestly distributed across a large fraction of the IT population (such as Fig 4a example unit 2).

Taken together, these results suggest that IT target match information is reflected by a weighted linear scheme and that target match performance depends on signals that are broadly distributed across most of the IT population.

### Projections along the IT linear decoding axis reflected behavioral confusions

Upon establishing that the format of IT target match information during the IDMS task was linear (on correct trials), we were interested in determining the degree to which behavioral confusions were reflected in the IT neural data. To measure this, we focused on the data recorded simultaneously across multiple units within each session, where all units observed the same errors. With this data, we trained the linear decoder to perform the same target match versus distractor classification described for Fig 5 using data from correct trials, and we measured cross-validated performance on pairs of condition-matched trials: one for which the monkey answered correctly, and the other for which the monkey made an error. On correct trials, target match performance grew with population size and reached above chance levels in populations of 24 units (Fig 6, black). On error trials, mean +/- SE of decoder performance fell below chance, and these results replicated across each monkey individually (Fig 6, white). These results establish that IT reflects behaviorally-relevant target match information insofar as projections of the IT population response along the FLD decoding axis co-vary with the monkeys’ behavior.

**Figure 6.**
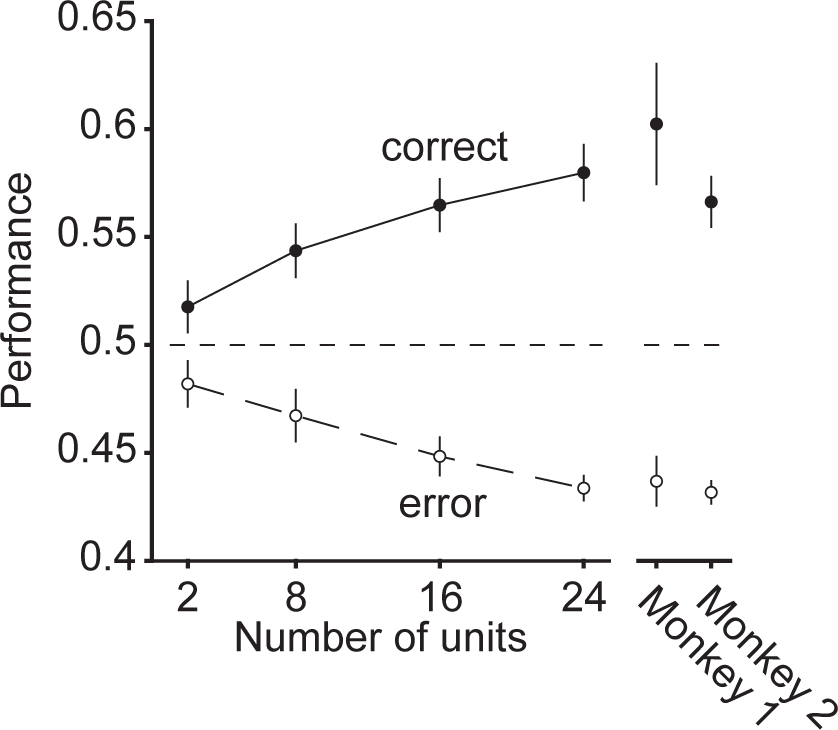
The IT FLD linear decoder axis reflects behavioral confusions. Linear decoder performance, applied to the simultaneously recorded data for each session, after training on correct trials and cross-validating on pairs of correct and error trials matched for condition. Error bars (SEM) reflect the variability that can be attributed to the random selection of units (for populations smaller than the full dataset) and the random assignment of training and testing trials in cross-validation. Results are shown for the data pooled across all sessions (main plot, n= 20 sessions) as well as when the sessions are parsed by those collected from each animal (monkey 1, n=10 sessions; monkey 2, n=10 sessions).

### Behaviorally-relevant target match signals were reflected as combinations of target match enhancement and suppression

As described in the introduction, the IT target match signal has largely been studied via the classic DMS paradigm (which includes the presentation of the cue at the beginning of the trial) and previous results have reported approximately balanced mixtures of target match enhancement and suppression (Miller and Desimone, 1994; Pagan et al., 2013). While some have speculated that target match enhancement alone reflects the behaviorally-relevant target match signal (Miller and Desimone, 1994), others have argued that enhancement and suppression are both behaviorally-relevant (Engel and Wang, 2011). The results presented above demonstrate that during the IDMS task, the representation of target match information is largely linear, and projections along the FLD weighted linear axis reflect behavioral confusions. To what degree does IT target match information, including the reflection of behavioral confusions, follow from units that reflect target match information with target enhancement (positive weights) as compared to target suppression (negative weights)? In our study, this question is of particular interest in light of the fact that our experimental design does not include the presentation of a cue at the beginning of each trial, and thus minimizes the degree to which target match suppression follows passively from stimulus repetition.

To investigate this question, we computed a target match modulation index for each unit as the average difference between the responses to the same images presented as target matches versus as distractors, divided by the sum of those two quantities. This index takes on positive values for target match enhancement and negative values for target match suppression. In both monkeys, this index was significantly shifted toward positive values (Fig 7a; Wilcoxon sign rank test, monkey 1: mean = 0.063 p = 8.44e^−6^; monkey 2: mean = 0.071, p = 2.11e^−7^). Notably, while these distributions were dominated by units that reflected target match enhancement, a small fraction of IT units in both monkeys reflected statistically reliable target match suppression as well (fraction of units that were significantly target match enhanced and suppressed, respectively, monkey 1: 49.1%, 17.6%; monkey 2: 41.7%, 8.3%; bootstrap significance test, p<0.01).

**Figure 7.**
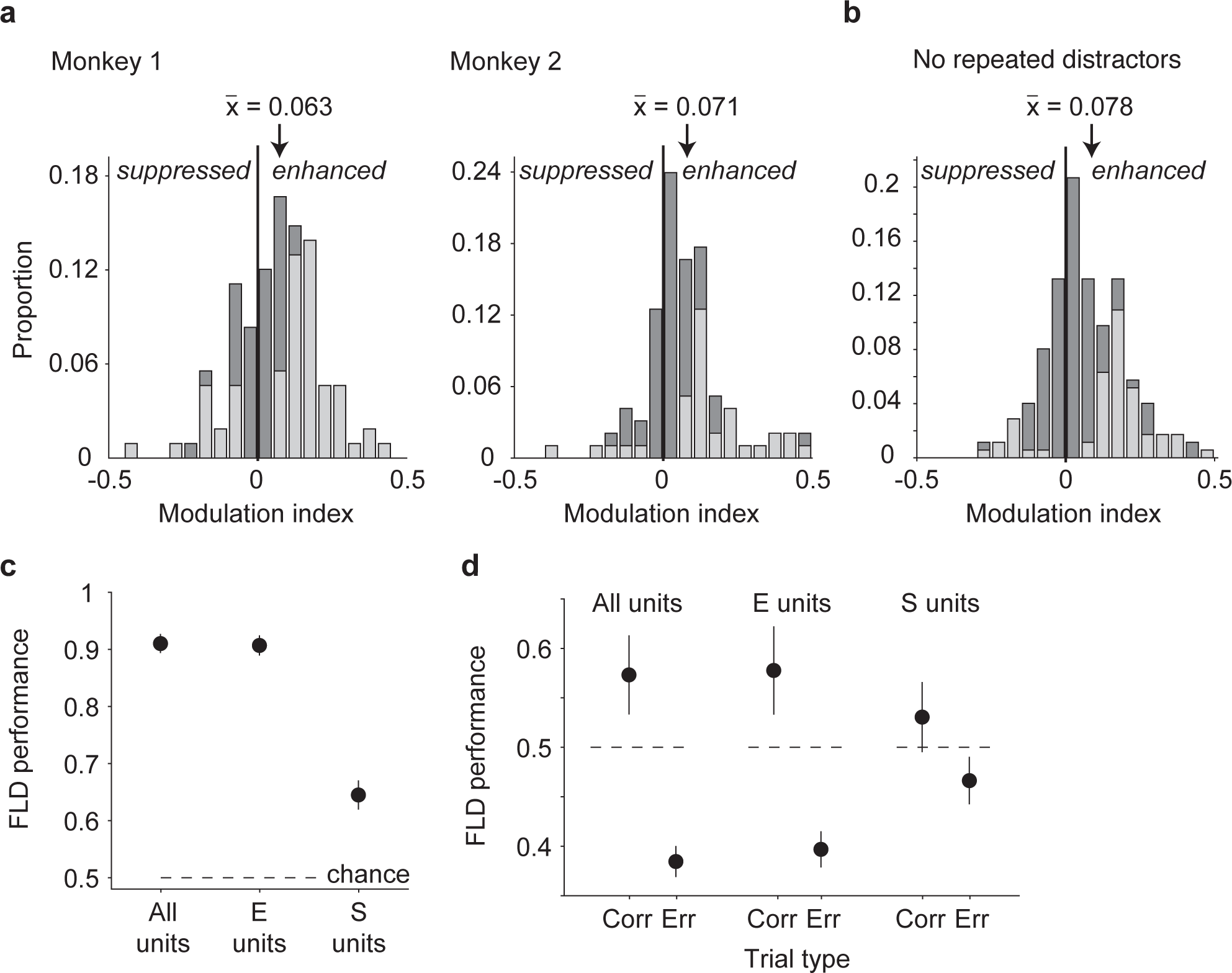
Target match signals are reflected as mixtures of enhancement and suppression. **a)** A target match modulation index, computed for each unit by calculating the mean spike count response to target matches and to distractors, and computing the ratio of the difference and the sum of these two values. Dark bars in each histogram indicate the proportions for all units (Monkey 1: n = 108; monkey 2: n = 96) whereas gray bars indicate the fractions of units whose responses to target matches versus distractors were statistically distinguishable (bootstrap significance test, p<0.01). Arrows indicate the distribution means. **b)** Target match modulation index, computed and plotted as in (a), but after excluding responses to repeated presentation of the same object within a trial. Included are units in which there were at least 10 repeated trials for each condition (n = 176 of 204 possible units). **c)** Performance of the FLD classifier for the combined population (n=204 units), computed for all units (as described for Fig 5b), target match enhanced units (“E units”) or target match suppressed units (“S units”). **d)** Performance of the FLD classifier for populations of size 24 recorded in each session when trained on correct trials and tested on condition-matched pairs of correct (“Corr.”) and error (“Err.”) trials (as described for Fig 6), computed for all units, E units, and S units.

In our experiment, the same images were not repeated within a trial but the same objects, presented under different transformations, could be. To what degree did the net target match enhancement that we observed follow from distractor suppression as a consequence of adaptation to object repetitions? To assess this, we recomputed target match modulation indices in a manner than only incorporated the responses to the first presentation of each object in a trial. Because this sub-selection reduced the number of distractor trials available for each condition, we equated these with equal numbers of (randomly selected) target match trials. A unit was only incorporated in the analysis if it had at least 10 trials per condition, yielding a subpopulation of 176 (of 204 possible) units. In the absence of distractor object repetitions, target match indices remained shifted toward net enhancement (Fig 7b; Wilcoxon sign rank test, mean = 0.078 p = 8.09e^−11^; fraction of units that were significantly target match enhanced and suppressed, respectively: 30.0%, 6.3%, bootstrap significance test, p <0.01), and the target match indices computed without repeated distractors were not statistically distinguishable from target match indices computed for the full dataset equated for numbers of trials, randomly selected (not shown; mean = 0.067, p = 0.33). We thus conclude that the dominance of target match enhancement in our population was not a consequence of distractor suppression that follows from object repetitions within a trial.

To determine the degree to which target match enhanced versus target match suppressed units contributed to population target match classification performance, we computed performance of the FLD linear decoder when isolated to the target match enhanced or target match suppressed subpopulations. More specifically, we focused on the combined data across the two monkeys (to maximize the numbers of units, particularly given small fraction that were target match suppressed), and we computed performance for variants of the FLD classifier in which the sign of modulation was computed for each unit based on the training data. Cross-validated performance was determined for either the subset of target match enhanced units or the subset of target match suppressed units with the goal of determining their respective contributions to overall population performance (while accounting for the fact that their proportions were not equal). When the analysis was isolated to target match enhanced units (“E units”), performance was virtually identical to the intact population (Fig 7c, mean+/- SEM performance for all units = 90.9+/-0.02% vs. E units = 90.6+/-0.02%), consistent with target match enhancement as the primary type of modulation driving population performance. When the analysis was isolated to target match suppressed units (“S units”), performance on correct trials was lower than that of the intact population but still well above chance (Fig 7c, performance for S units = 64.4+/0.03%). This suggests that while target match suppressed units are smaller in number, the target match suppressed units that do exist do in fact carry reliable target match signals.

What were the relative contributions of E units versus S units to error trial confusions? To address this question, we repeated the error trial analysis described above for Figure 6, but isolated to E or S units. Specifically, we repeated the analysis presented in Figure 6 where we considered the simultaneously recorded data collected across 24 units for each session, but isolated to the E or S units as described for Figure 7c (based on the training data), and we compared cross-validated performance on condition-matched correct versus error trials. E units classified correct trials above chance and misclassified error trials below chance at rates similar to the entire population (Fig 7d, “All units” vs. “E units”), consistent with a larger overall proportion of E units. In contrast, performance of the S units on correct trials was weaker and mean +/- SEM performance was not above chance (53.0+/- 0.04%; Fig 7d “S units, Corr.”), consistent with smaller numbers of these units in IT. Similarly, performance of S units on correct trials was slightly but not significantly higher than performance on error trials (mean +/- SE performance on error trials = 46.6+/-0.02%; p = 0.090, Fig 7d, “S units, Err.”). These results indicate that the reflection of behavioral confusions in the IT neural data arises primarily from the activity of E units, but suggest that behavioral confusions may also be weakly reflected in S units.

As a complementary analysis of behavioral relevance, we also examined the degree to which the responses to target matches reflected pre-saccadic activity by comparing the same responses time-locked to stimulus onset versus saccade onset (Fig 8). The saccade-aligned response was smaller and more diffuse than the stimulus-aligned response and saccade-aligned responses peaked well before saccade onset (~200 ms), suggesting that on average, IT responses to target matches do not reflect characteristic pre-saccadic activity.

**Figure 8.**
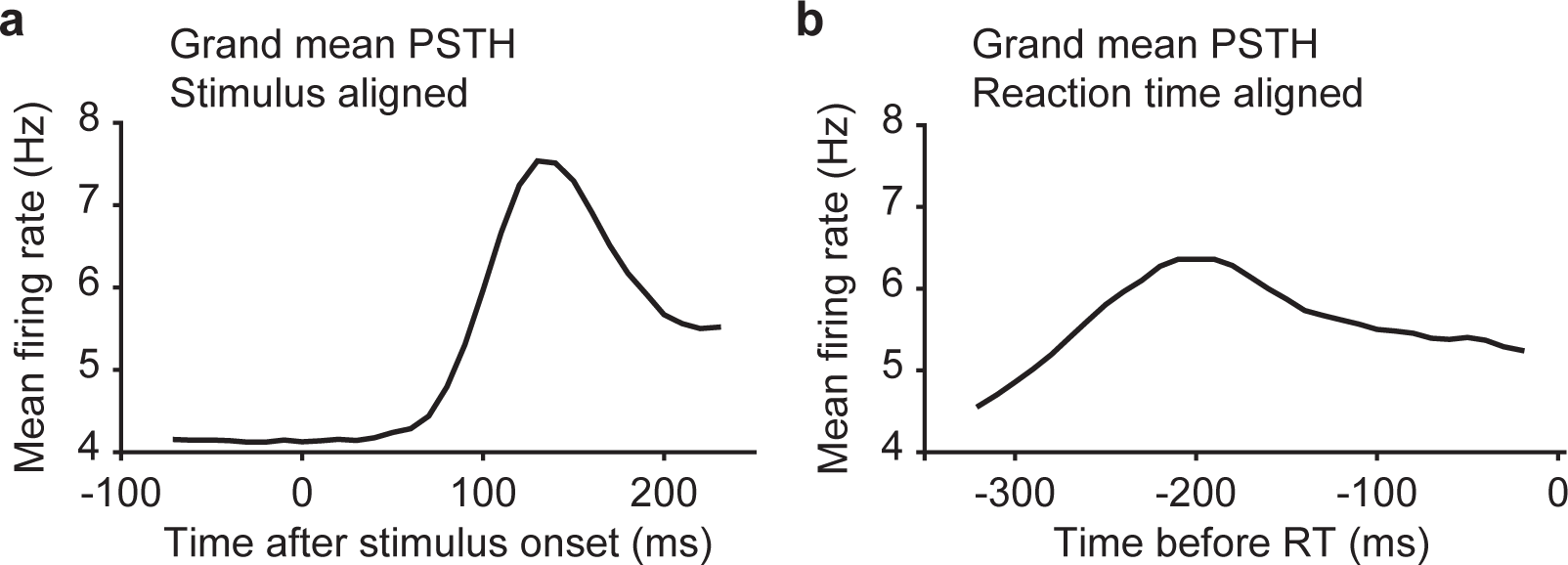
Comparison of stimulus-aligned versus reaction time-aligned responses to target matches. **a)** Grand mean PSTH across all units (n=204) for all target match stimuli, aligned to stimulus onset. **b)** Grand mean PSTH across all units (n=204) for all target match stimuli, aligned to behavioral reaction time.

Together, these results suggest that in the IDMS experiment, target match signals were dominated by target match enhancement, but a smaller, target match suppressed subpopulation exists as well. Additionally, they suggest that the reflection of behavioral confusions in IT neural responses could largely be attributed to units that are target match enhanced, but behavioral confusions were weakly reflected in units that are target match suppressed. Finally, while IT responses reflect behavioral confusions, they were not well-aligned to reaction times.

### The IT target match representation was configured to minimize interference with IT visual represen ta tions

As a final topic of interest, we wanted to understand how the representation of target match information was multiplexed with visual representations in IT and more specifically, whether IT had a means of minimizing the potentially detrimental impact of mixing these two types of signals. One possible way to achieve this is multiplicative rescaling, as described in Figure 1. To what degree is this happening in IT? As a first step toward addressing this question, we quantified the impact of target match modulation as the representational similarity between the IT population response vectors corresponding to the same images presented as target matches versus as distractors, using a scale-invariant measure of similarity (the Pearson correlation, reviewed by Kriegeskorte and Kievit, 2013). More specifically, we measured the Pearson correlation between pairs of population response vectors via a split-halves procedure (see Methods), and we compared the representational similarity for the same images presented as target matches versus as distractors with other benchmarks in our experiment, including: within the same experimental condition (i.e. random splits across repeated trials); between images containing different transformations of the same object; and between images containing different objects.

Shown in Figure 9a is the representational similarity matrix corresponding to all possible pairwise combinations of the 20 images used in this experiment, averaged across the matrices computed when the pairs of response vectors under consideration were target matches and when they were distractors, computed with spike count windows 80-250 ms relative to stimulus onset (see Methods). The matrix is organized such that the five transformations corresponding to each object are grouped together. Figure 9b reorganizes the data into plots of the mean and standard error of representational similarity computed for different pairwise comparisons. As expected, we found that the representational similarity was the highest for random splits of the trials corresponding to the same images, presented under the same conditions (Fig 9b, “Same image & condition”, mean = 0.43), which can be regarded as the noise ceiling in our data. In comparison, the representational similarity was significantly lower for different transformations of the same object (Fig 9b, “Different transforms.”; mean = 0.14; p = 1.14e^−8^) as well as for different objects (Fig 9b, “Different objects”; mean = −0.02; p = 1.92e^−29^). We note that a representational similarity value of zero reflects the benchmark of IT population responses that are orthogonal, and this was the case for the representation of different objects in IT. It was also the case that representational similarity was significantly lower for different objects as compared to different transformations of the same object (p=1.43e^−7^), consistent with an IT representation that was tolerant to changes in identity-preserving transformations. With these benchmarks established, what impact did target match modulation have on IT visual representations? The average representational similarity for the same images presented as target matches as compared to distractors was significantly lower than the noise ceiling (Fig 9b, “Matches versus distractors”; mean = 0.28; p = 2.09e^−7^) but was significantly higher than presenting the same object under a new transformation (Fig 9b, p = 0.0016) or presenting a different object (Fig 9b, p = 3.057e^−20^). These results suggest that the multiplexing of IT target match signals was not perfect, but also had a smaller impact on the population response than changing either the transformation in which an object was viewed in or the object in view. These results, computed for broad spike count windows (80-250 ms), were qualitatively replicated in narrower windows positioned early (80-130 ms), midway (140-190 ms) and late (200-250 ms) relative to stimulus onset (Fig 9c). Most notably, representational similarity for matches and distractors remained significantly higher than representational similarity for different transformations of the same object in all epochs (Fig 9c, “Mtch. v. Dstr.” vs. “Diff. trans.”, early p = 0.0023, mid p = 0.0081, late p = 0.0092). These results confirm that the impact of target match modulation on IT population representational similarity remains modest throughout the stimulus-evoked response period.

**Figure 9.**
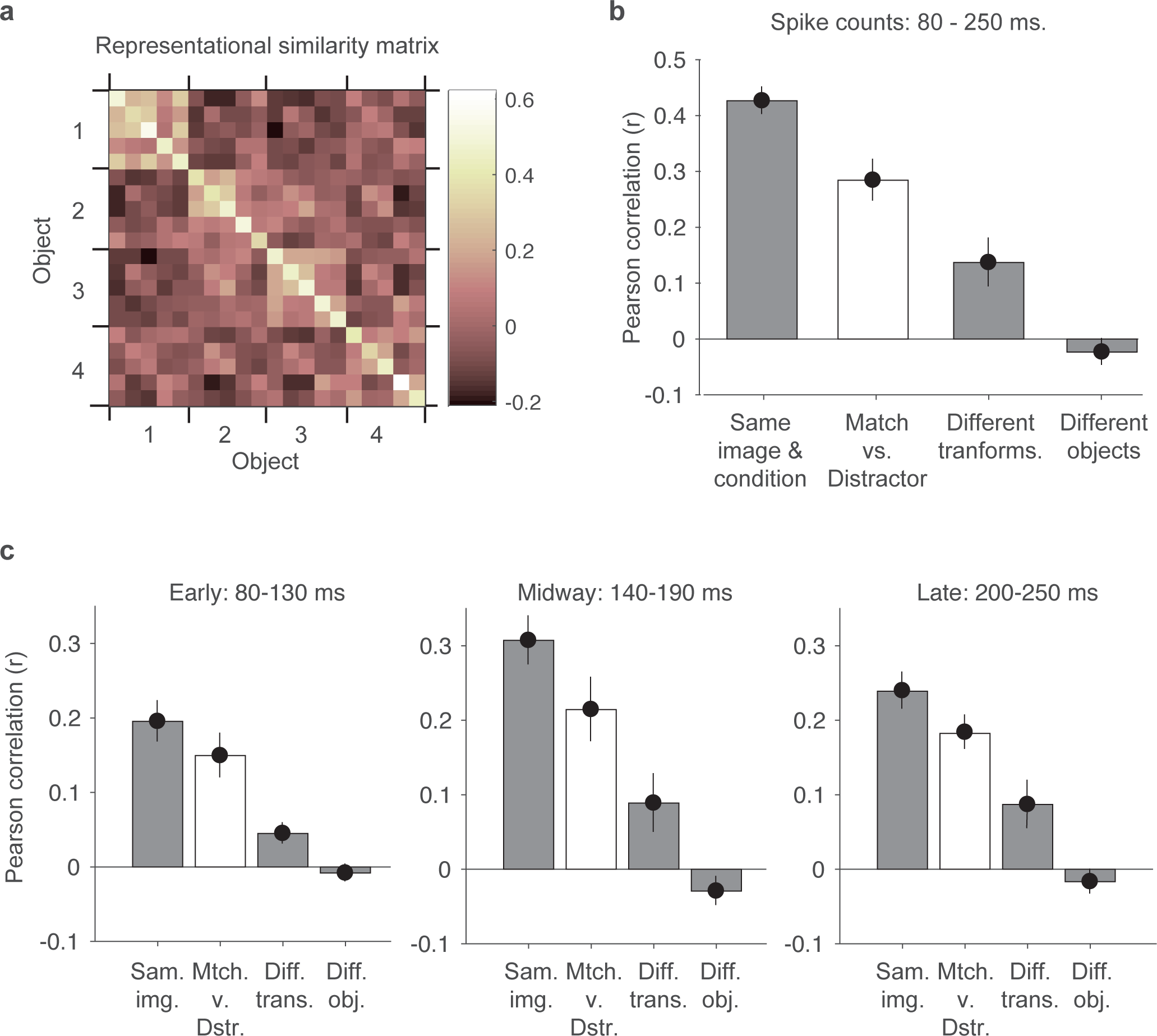
Target match signaling has minimal impact on the IT visual population response. **a)** The representational similarity matrix, computed as the average Pearson correlation between the population response vectors computed for all possible pairs of images. Before computing the correlations between pairs of population response vectors, the responses of each unit were z-normalized to ensure that correlation values were not impacted by differences in overall firing rates across units (see Methods). Correlations were computed based on a split halves procedure. Shown are the average correlations, computed between images with a fixed target and averaged across all possible targets, as well as averaged across 1000 random splits. The matrix is organized such that different transformations of the same object are grouped together, in the same order as depicted in Fig 2. **b)** The average representational similarity, computed across: “Same image and condition”: different random splits of the 20 trials into two sets of 10 trials each; “Different transforms.”: images containing different transformations of the same object, computed with a fixed target identity; “Different objects”: images containing different objects, computed with a fixed target identity; “Match versus distractor”: the same image viewed as a target match as compared to as a distractor, averaged across all 9 possible distractor combinations (see Methods). **c)** The analysis described for panel b applied to different time epochs. Error bars (SEM) reflect variability across the 20 images.

To what degree does the modest impact of target match modulation follow from the multiplicative mechanism highlighted in Figure 1? One requirement for multiplicative population responses are individual units whose responses are themselves multiplicatively rescaled. To determine the degree to which our recorded IT units were multiplicative, we computed the impact of target match modulation as a function of stimulus rank and compared it to the benchmarks expected for multiplicative rescaling as well as other alternatives (including subtraction and sharpening; Fig 10a,c). Specifically, we ranked the responses of each unit to the 20 images separately (after averaging across target matches and distractors), and we then computed the average across all units at each rank for target matches and distractors separately. Average IT target match modulation was much better described as multiplicative than as subtractive or sharpening (Fig 10b,d).

**Figure 10.**
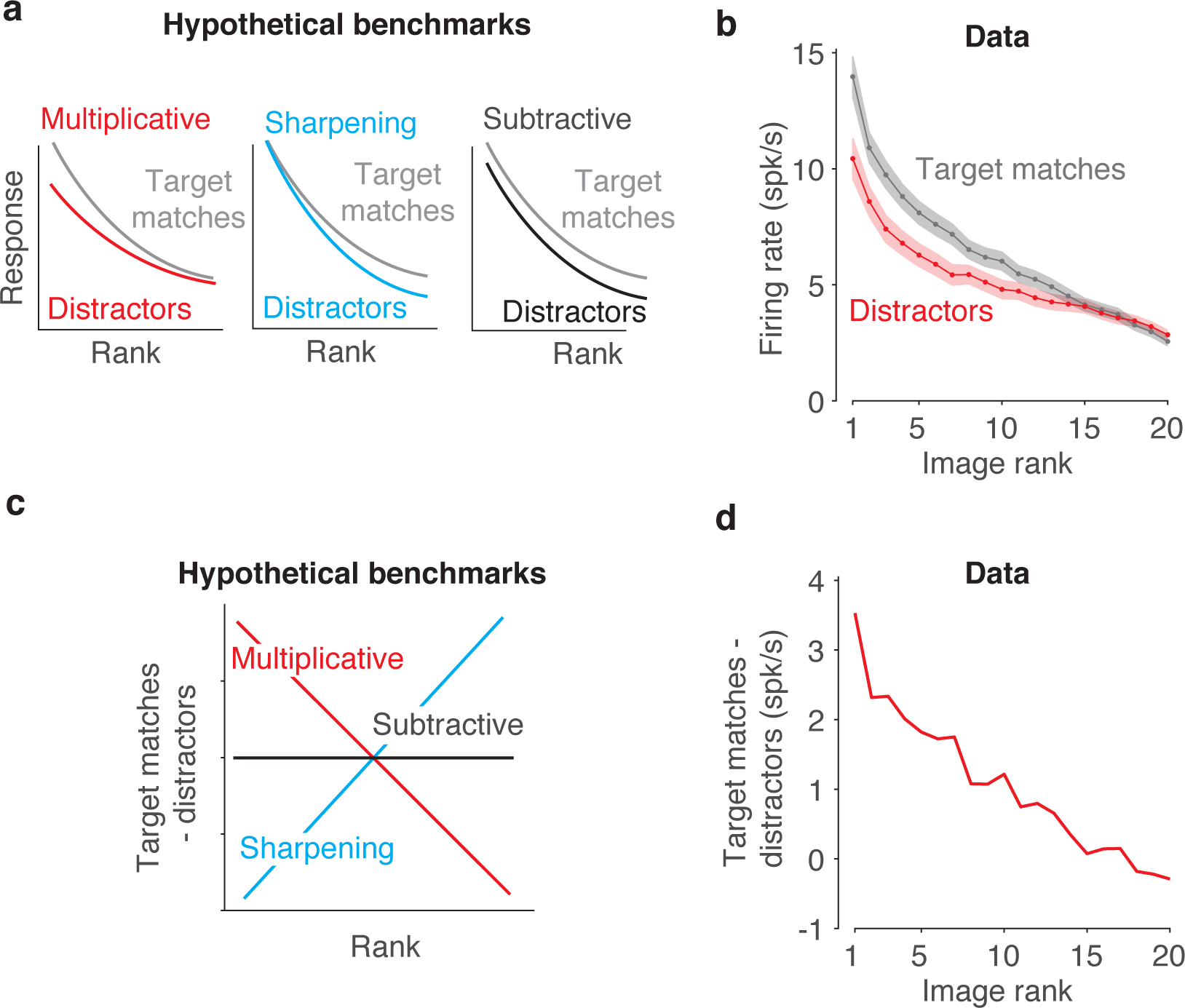
The impact of target match modulation on the visual responses of individual units. **a)** Cartoon depiction of the impact of different types of target match modulation on the rank-ordered responses to different images. **b)** Mean and SEM of the rank-order responses across units, after ranking the responses for each unit separately (based on the averaged response to target matches and distractors). **c)** The cartoons in panel a, replotted as the difference between target matches and distractors at each rank to visualize the differences between them. **d)** The analysis described in panel c, applied to the data in panel b, reveals that the impact of target match modulation is better described as multiplicative than as subtractive or as sharpening.

A second requirement for multiplicative population response vectors is homogeneity in target match modulation across units (Fig 11a, cyan). Variation across units in terms of the magnitudes of target match modulation (Fig 11a, left, red), and/or variation that includes mixtures of target match enhancement and suppression (Fig 11a, right, red) can produce changes in population response vector positions that could be confounded with changes in the visual identity, if the variations were sufficiently large. Where does the amount of target match modulation heterogeneity that we observed (e.g. Fig 7a) fall relative to the benchmarks of the best versus worst format that it could possibly take? To investigate this question, we performed a series of data-based simulations targeted at benchmarking our results relative to “best case” and “worse case” scenarios for multiplexing given the magnitudes of target match modulation in our data. As a first “replication” simulation, we replicated the responses recorded for each unit by preserving the magnitudes and types of signals as well as each unit’s grand mean spike count and we simulated trial variability with an independent, Poisson process (see Methods). The pattern of representational similarities reflected in the raw data (Fig 9b) were approximated in simulation (Fig 11b), suggesting that this simulation procedure was effective at capturing important elements of the data. In the other simulations described below, we began in the same way: by preserving the amounts and types of visual, target and residual modulation recorded in each unit, as well as each unit’s grand mean firing rate. What differed between the simulations was how that target match modulation was distributed across units.

To simulate the “best case scenario” in our data, we approximated multiplicative rescaling by distributing the total target match modulation across units in equal proportions relative to their magnitudes of visual modulation. In this simulation, target match modulation was introduced with the same sign (target match enhancement) across all units, consistent with the average sign reflected in the raw data (Fig 7a). Representational similarity between target matches and distractors in this multiplicative, same-sign simulation was statistically indistinguishable from the noise ceiling (Fig 11c, p = 0.395), confirming intuitions that a population can (in principle) multiplex target match signals in a multiplicative manner that has minimal interference with visual representations. To simulate a “worse case scenario” for our data, we increased the amount of target match modulation heterogeneity across units by both distributing target match modulation uniformly (as opposed to proportionally) across units as well as preserving the original sign of each unit’s target match modulation (i.e. target match enhancement or suppression). Representational similarity between target matches and distractors in this uniform, mixed-sign simulation fell to levels measured for different transformations of the same object (Fig 11c), confirming that our data do not reflect a “worst case scenario” given the magnitudes of target match modulation that we observed. Together, these results suggest that in line with Fig 1, the impact of target match modulation on IT visual representations is modest (Fig 9) as a consequence of modulation that is approximately (albeit imperfectly) multiplicative, due both to individual units with target match modulation that is multiplicative on average, as well as target match modulation that is approximately (albeit imperfectly) functionally homogenous.

**Figure 11.**
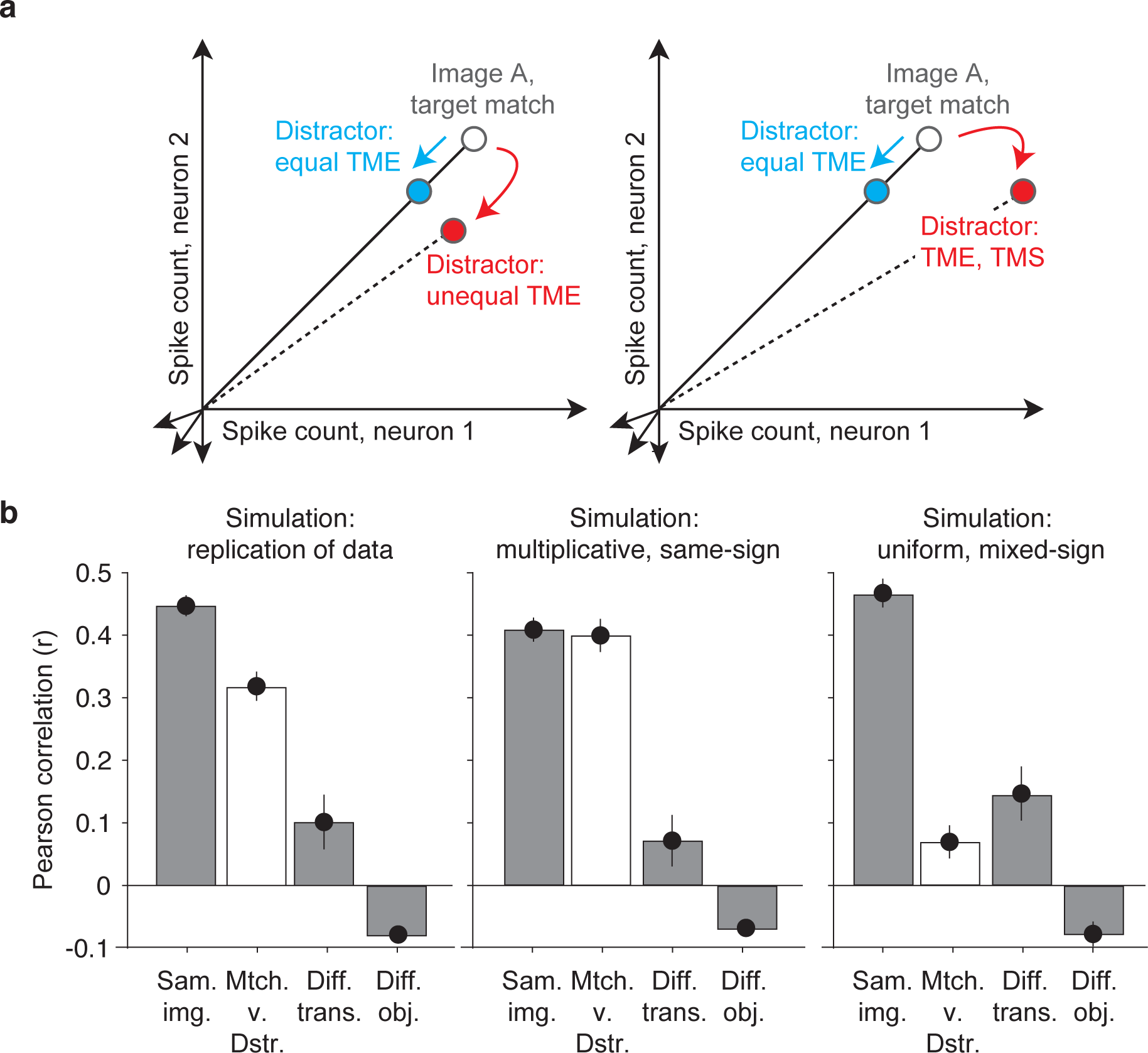
Benchmarking the impact of target match modulation heterogeneity across units. **a)** Cartoon depiction of how heterogeneity across units in target match modulation magnitudes (left) and modulation signs (right) can lead to changes in the population response to the same images presented as target matches versus distractors. **b)** Three simulated variants of the recorded data (see Results), including target match modulation for each unit that was: replicated; enforced to be multiplicative and reflected with the same-sign across all units (i.e. target match enhancement); enforced to be uniform and reflected with mixed-signs across units (i.e. target match enhancement or suppression, as determined by the original data).

## Discussion

Successfully finding a sought target object, such as your car keys, requires your brain to compute a target match signal that reports when a target is in view. Target match signals have been reported to exist in IT, but these signals are not well understood, particularly in the context of the real-world problem of searching for an object that can appear at different identity-preserving transformations. We recorded responses in IT as two monkeys performed a delayed-match-to-sample task in which a target object could appear at different positions, sizes, and background contexts. We found that the IT population reflected a target match representation that was largely linear, and that it reflected behavioral confusions on trials in which the monkeys made errors. IT target match signals were broadly distributed across most IT units, and while they were dominated by target match enhancement, we also found evidence for reliable target match suppression. Finally, we found that IT target match modulation was configured in such a manner as to minimally impact IT visual representations. Together, these results support the existence of a robust, behaviorally-relevant target match representation in IT that is multiplexed with IT visual representations.

Our results support the existence of a robust target match representation in IT during this task that reflects confusions on trials in which the monkeys make errors (Fig 6); this result has not been reported previously. One earlier study also explored the responses of IT neurons in the context of a DMS task in which, like ours, the objects could appear at different identity-preserving transformations (Leuschow et al., 1994), but this study did not sort neural responses based on behavior. Another study examined IT neural responses as monkeys performed a visual target search task that involved free viewing as well as image manipulation during the time of the saccade (Mruczek and Sheinberg, 2007). They reported higher firing rates in IT neurons during trial sequence that normally led to a reward (an association between a target object and a saccade to a response target) versus swap trials in which this sequence was disrupted. Another study (from our lab) used a classic DMS design reported that IT population classifications on error trials fell to chance (Pagan et al., 2013), but this study did not find evidence for significant error trial misclassifications.

IT target match signals have been investigated most extensively in IT via a classic version of the delayed-match-to-sample (DMS) paradigm where each trial begins with a visual cue indicating the identity of the target object, and this cue is often the same image as the target match (Eskandar et al., 1992; Miller and Desimone, 1994; Pagan et al., 2013). In this paradigm, approximately half of all IT neurons that differentiate target matches from distractors do so with enhanced responses to matches whereas the other half are match suppressed (Miller and Desimone, 1994; Pagan et al., 2013). Because match suppressed responses also follow from the repetition of distractors within a trial, some have speculated that the match enhanced neurons alone carry behaviorally-relevant target match information (Miller and Desimone, 1994). In general agreement with those notions, the target match signal is dominated by target match enhancement in situations where the cue and target match are presented at different locations (Chelazzi et al., 1993). Conversely, others have argued that a representation comprised exclusively of match enhanced neurons would confuse the presence of a match with modulations that evoke changes in overall firing rate, such as changes in stimulus contrast (Engel and Wang, 2011). Additionally, these authors proposed that match suppressed neurons could be used in these cases to disambiguate target match versus stimulus-induced modulation. In our experiment, the IDMS task was run in blocks containing a fixed target to minimize the impact of passive stimulus repetition of the target match. We found evidence for net target match enhancement in our data (Fig 7a), and that this in turn translated into a type of homogeneity that minimized the potentially detrimental impact of target match modulation on visual representations (Fig 11). However, we also found evidence for a smaller subpopulation of units that reflected reliable target match suppression. Whether the amount of target match suppression that we observed is sufficient for the disambiguation strategy proposed by Engel and Wang (2011) is thus unclear - because our experiment did not include variation in parameters that change overall firing rate (such as contrast), we cannot directly test it with our data.

How does the target match signal arrive in IT? Computation of the target match signal requires a comparison of the content of the currently-viewed scene with a remembered representation of the sought target. The existence of target match signals in IT could reflect the implementation of the comparison in IT itself or, alternatively, this comparison might be implemented in a higher-order brain area (such as prefrontal cortex) and fed-back to IT. Examination of the timing of the arrival of this signal in IT (which peaks at 150 ms; Fig 4b) relative to the monkeys’ median reaction times (~340 ms; Fig 2e), does not rule out the former scenario. The fact that neural responses to target matches were more time locked to stimulus onset than they are to reaction times suggests that this activity does not reflect classic signatures of motor preparation. Additional insights into whether or not target match signals are computed in IT might be gained through analyses of the responses on cue trials, particularly with regard to whether signatures of the visually-evoked responses to cues persist throughout each block, however, our experimental design included too few cue presentations for such analyses. Thus while our data are consistent with target match computations within IT cortex, we cannot definitively distinguish this proposal from alternative scenarios with this data. Additionally, in this study monkeys were trained extensively on the images used in these experiments and future experiments will be required to address the degree to which these results hold under more everyday conditions in which monkeys are viewing images and objects for the first time.

In a previous series of reports, we investigated target match signals in the context of the classic DMS design in which target matches were repeats of cues presented earlier in the trial and each object was presented on a gray background (Pagan and Rust, 2014a; Pagan et al., 2016; Pagan et al., 2013). One of our main findings from that work was that the IT target match representation was reflected in a partially nonlinearly separable format, whereas an IT downstream projection area, perirhinal cortex, contained the same amount of target match information but in a format that was largely linearly separable. In the data we present here, we did not find evidence for a nonlinear component of the IT target match representation, reflected as consistently higher performance of a maximum likelihood as compared to linear decoder (Fig 5b). The source of these differences is unclear. They could arise from the fact that the IDMS task requires an “invariant” visual representation of object identity, which first emerges in a linearly separable format in the brain area that we are recording from (Rust and DiCarlo, 2010), whereas in more classic forms of the DMS task, the integration of visual and target information could happen in a different manner and/or a different brain area. Alternatively, these differences could arise from the fact that during IDMS, images are not repeated within a trial, and the stronger nonlinear component revealed in DMS may be produced by stimulus repetition. It may also be the case that nonlinearly separable information is in fact present in IT during IDMS but was not detectable under the specific conditions used in our experiments. For example, the proportion of nonlinearly separable information grows as a function of population size, and it may be the case that it is detectable during IDMS for larger sized populations. Our current data cannot distinguish between these alternatives.

Our results also add to the growing literature that suggests the brain “mixes” the modulations for different task-relevant parameters within individual neurons, even at the highest stages of processing (Freedman and Assad, 2009; Kobak et al., 2016; Mante et al., 2013; Meister et al., 2013; Raposo et al., 2014; Rigotti et al., 2013; Rishel et al., 2013; Zoccolan et al., 2007). A number of explanations have been proposed to account for mixed selectivity. Some studies have documented situations in which signal mixing is an inevitable consequence of the computations required for certain tasks, such as identifying objects invariant to the view in which they appear (Zoccolan et al., 2007). Others have suggested that mixed selectivity may be an essential component of the substrate required to maintain a representation that can rapidly and flexibly switch with changing task demands (Raposo et al., 2014; Rigotti et al., 2013). Still others have maintained that broad tuning across different types of parameters is important for learning new associations (Barak et al., 2013). Our results suggest that IT mixes visual and target match information within individual units. This could reflect the fact that the comparison of visual and target match information happens within IT itself, and multiplexing is simply a byproduct of that computation. Alternatively, if the comparison is performed elsewhere, this would reflect its feedback to IT for some unknown purpose. In either case, our results suggest that the multiplexing happens in a manner that is largely but imperfectly multiplicative (Fig 10-11) and thus configured to minimize interference of visual representations when also signaling target match information.

## Acknowledgements

We thank Margot P. Wohl for her contributions to early phases of this work. This work was supported by the National Eye Institute of the National Institutes of Health (award R01EY020851), the Simons Foundation (through an award from the Simons Collaboration on the Global Brain), and the McKnight Endowment for Neuroscience.

## METHODS

Experiments were performed on two adult male rhesus macaque monkeys *(Macaca mulatta)* with implanted head posts and recording chambers. All procedures were performed in accordance with the guidelines of the University of Pennsylvania Institutional Animal Care and Use Committee and this study was approved under protocol 804222.

### The invariant delayed-match-to-sample (IDMS) task

All behavioral training and testing was performed using standard operant conditioning (juice reward), head stabilization, and high-accuracy, infrared video eye tracking. Stimuli were presented on an LCD monitor with an 85 Hz refresh rate using customized software (http://mworks-project.org).

As an overview, the monkeys’ task required an eye movement response to a specific location when a target object appeared within a sequence of distractor images (Fig 2a). Objects were presented across variation in the objects’ position, size and background context (Fig 2b). Monkeys viewed a fixed set of 20 images across switches in the identity of 4 target objects, each presented at 5 identity-preserving transformations (Fig 2c). Monkeys were trained extensively on the set of 20 images shown in Fig 2b before testing. We ran the task in short blocks (~3 min) with a fixed target before another target was pseudorandomly selected. Our design included two types of trials: cue trials and test trials (Fig 2a). Only test trials were analyzed for this report.

Trials were initiated by the monkey fixating on a red dot (0.15°) in the center of a gray screen, within a square window of ±1.5°, followed by a 250 ms delay before a stimulus appeared. Cue trials, which indicated the current target object, were presented at the beginning of each block and after three subsequent trials with incorrect responses. To minimize confusion, cue trials were designed to be distinct from test trials and began with the presentation of an image of each object that was distinct from the images used on test trials (a large version of the object presented at the center of gaze on a gray background; Fig 2a). Test trials, which are the focus of this report, always began with a distractor image, and neural responses to this image were discarded to minimize non-stationarities such as stimulus onset effects. Distractors were drawn randomly from a pool of 15 possible images within each block without replacement until each distractor was presented once on a correct trial, and the images were then re-randomized. On most trials, a random number of 1-6 distractors were presented, followed by a target match (Fig 2a). On a small fraction of trials, 7 distractors were shown, and the monkey was rewarded for fixating through all distractors. Each stimulus was presented for 400 ms (or until the monkeys’ eyes left the fixation window) and was immediately followed by the presentation of the next stimulus. Following the onset of a target match image, monkeys were rewarded for making a saccade to a response target within a window of 75 - 600 ms to receive a juice reward. In monkey 1 this target was positioned 10 degrees above fixation; in monkey 2 it was 10 degrees below fixation. If 400 ms following target onset had elapsed and the monkey had not moved its eyes, a distractor stimulus was immediately presented. If the monkey continued fixating beyond the required reaction time, the trial was considered a “miss”. False alarms were differentiated from fixation breaks via a comparison of the monkeys’ eye movements with the characteristic pattern of eye movements on correct trials: false alarms were characterized by the eyes leaving the fixation window via its top (monkey 1) or bottom (monkey 2) outside the allowable correct response period and traveling more than 0.5 degrees whereas fixation breaks were characterized by the eyes leaving the fixation window in any other way. Within each block, 4 repeated presentations of the 20 images were collected, and a new target object was then pseudorandomly selected. Following the presentation of all 4 objects as targets, the targets were re-randomized. At least 20 repeats of each condition were collected. Overall, monkeys performed this task with high accuracy. Disregarding fixation breaks (monkey 1: 11% of trials, monkey 2: 8% of trials), percent correct on the remaining trials was as follows: monkey 1: 96% correct, 1% false alarms, and 3% misses; monkey 2: 87% correct, 3% false alarms, and 10% misses.

### Neural recording

The activity of neurons in IT was recorded via a single recording chamber in each monkey. Chamber placement was guided by anatomical magnetic resonance images in both monkeys, and in one monkey, Brainsight neuronavigation (https://www.rogue-research.com/). The region of IT recorded was located on the ventral surface of the brain, over an area that spanned 4 mm lateral to the anterior middle temporal sulcus and 15-19 mm anterior to the ear canals. Neural activity was largely recorded with 24-channel U probes (Plexon, Inc) with linearly arranged recording sites spaced with 100 mm intervals, with a handful of units recorded with single electrodes (Alpha Omega, glass-coated tungsten). Continuous, wideband neural signals were amplified, digitized at 40 kHz and stored using the OmniPlex Data Acquisition System (Plexon). Spike sorting was done manually offline (Plexon Offline Sorter). At least one candidate unit was identified on each recording channel, and 2-3 units were occasionally identified on the same channel. Spike sorting was performed blind to any experimental conditions to avoid bias. A multi-channel recording session was included in the analysis if the animal performed the task until the completion of 20 correct trials per stimulus condition, there was no external noise source confounding the detection of spike waveforms, and the session included a threshold number of task modulated units (>4 on 24 channels). The sample size (number of units recorded) was chosen to approximately match our previous work (Pagan and Rust, 2014a; Pagan et al., 2016; Pagan et al., 2013).

For all the analyses presented in this paper except Fig 4b,d, Fig 8, and Fig 9c, we measured neural responses by counting spikes in a window that began 80 ms after stimulus onset and ended at 250 ms. On 1.9% of all correct target match presentations, the monkeys had reaction times faster than 250 ms, and those instances were excluded from analysis such that spikes were only counted during periods of fixation. When combining the units recorded across sessions into a larger pseudopopulation, we screened for units that met three criteria. First, units had to be modulated by our task, as quantified by a one-way ANOVA applied to our neural responses (80 conditions * 20 repeats) with p < 0.01. Second, we applied a loose criterion on recording stability, as quantified by calculating the variance-to-mean for each unit (computed by fitting the relationship between the mean and variance of spike count across the 80 conditions), and eliminating units with a variance-to-mean ratio > 5. Finally, we applied a loose criterion on unit recording isolation, quantified by calculating the signal-to-noise ratio (SNR) of the waveform (as the difference between the maximum and minimum points of the average waveform, divided by twice the standard deviation across the differences between each waveform and the mean waveform), and excluding (multi)units with an SNR < 2. This yielded a pseudopopulation of 204 units (of 563 possible units), including 108 units from monkey 1 and 96 units from monkey 2.

### Quantifying single-unit modulation magnitudes

To quantify the degree to which individual units were modulated by different types of task parameters (Fig 4b-d), we applied a bias-corrected procedure described in detail by (Pagan and Rust, 2014b) and summarized here. Our measure of modulation is similar to a multi-way ANOVA, with important extensions. Specifically, a two-way ANOVA applied to a unit’s responses (configured into a matrix of 4 targets * 20 images * 20 trials for each condition) would parse the total response variance into two linear terms, a nonlinear interaction term, and an error term. We make 3 extensions to the ANOVA analysis. First, an ANOVA returns measures of variance (in units of spike counts squared) whereas we compute measures of standard deviation (in units of spike count) such that our measures of modulation are intuitive (e.g., doubling firing rates causes signals to double as opposed to quadruple). Second, while the linear terms of the ANOVA map onto our “visual” and “target identity” modulations (after squaring), we split the ANOVA nonlinear interaction term into two terms, including target match modulation (i.e. Fig 2c gray versus white) and all other nonlinear “residual” modulation. This parsing is essential, as target match modulation corresponds to the signal for the IDMS task whereas the other types of modulations are not. Finally, raw ANOVA values are biased by trial-by-trial variability (which the ANOVA addresses by computing the probability that each term is higher than chance given this noise) whereas our measures of modulation are bias-corrected to provide an unbiased estimate of modulation magnitude.

The procedure begins by developing an orthonormal basis of 80 vectors designed to capture all types of modulation with intuitive groupings. The number of each type is imposed by the experimental design. This basis ***b*** included vectors ***b*_*t*_** that reflected 1) the grand mean spike count across all conditions ***(b*_*t*_**, 1 dimension), 2) whether the object in view was a target or a distractor ***(b*_*2*_**, 1 dimension), 3) visual image identity ***(b_3_ - b_21_***, 19 dimensions), 4) target object identity ***(b_22_ - b_2A_***, 3 dimensions), and 5) “residual”, nonlinear interactions between target and object identity not captured by target match modulation ***(b_25_ - b_80_***, 56 dimensions). A Gram-Schmidt process was used to convert an initially designed set of vectors into an orthonormal basis.

Because this basis spans the space of all possible responses for our task, each trial-averaged vector of spike count responses to the 80 experimental conditions ***R*** can be re-expressed as a weighted sum of these basis vectors. To quantify the amounts of each type of modulation reflected by each unit, we began by computing the squared projection of each basis vector ***b*_*t*_** and ***R.*** An analytical bias correction, described and verified in (Pagan and Rust, 2014b), was then subtracted from this value:

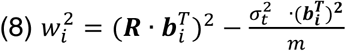

where 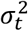 indicates the trial variance, averaged across conditions (n=80), and where m indicates the number of trials (m=20). When more than one dimension existed for a type of modulation, we summed values of the same type. Next, we applied a normalization factor (1/(n-1)) to convert these summed values into variances. Finally, we computed the square root of these quantities to convert them into modulation measures that reflected the number of spike count standard deviations around each unit’s grand mean spike count.

Target match modulation was thus computed as:

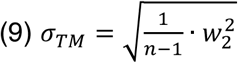

visual modulation was computed as:

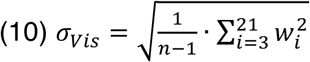

target identity modulation was computed as:

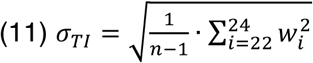

and residual modulation was computed as:

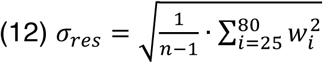

When estimating modulation population means (Fig 4b,c), the bias-corrected squared values were averaged across units before taking the square root. Because these measures were not normally distributed, standard error about the mean was computed via a bootstrap procedure. On each iteration of the bootstrap (across 1000 iterations), we randomly sampled values from the modulation values for each unit in the population, with replacement. Standard error was computed as the standard deviation across the means of these newly created populations.

To quantify the sign of the modulation corresponding to whether an image was presented as a target match versus as a distractor (Fig 7a,b), we calculated a target match modulation index for each unit by computing its mean spike count response to target matches and to distractors, and computing the ratio of their difference and their sum.

### Population performance

To determine the performance of the IT population at classifying target matches versus distractors, we applied two types of decoders: a Fisher Linear Discriminant (a linear decoder) and Maximum Likelihood decoder (a nonlinear decoder) using approaches that are described previously in detail (Pagan et al., 2013) and are summarized here.

When applied to the pseudopopulation data (Fig 5b, Fig 7b), all decoders were cross-validated with the same resampling procedure. On each iteration of the resampling, we randomly shuffled the trials for each condition and for each unit, and (for numbers of units less than the full population size) randomly selected units. On each iteration, 18 trials from each condition were used for training the decoder, 1 trial was used to determine a value for regularization, and 1 trial from each condition was used for cross-validated measurement of performance.

To ensure that decoder performance was not biased by unequal numbers of target matches and distractors, on each iteration of the resampling we included 20 target match conditions and 20 (of 60 possible) distractor conditions. Each set of 20 distractors was selected to span all possible combinations of mismatched object and target identities (e.g. objects 1, 2, 3, 4 paired with targets 4, 3, 2, 1), of which there are 9 possible sets. To compute proportion correct, a mean performance value was computed on each resampling iteration by averaging binary performance outcomes across the 9 possible sets of target matches and distractors, each which contained 40 test trials. Mean and standard error of performance was computed as the mean and standard deviation of performance across 1000 resampling iterations. Standard error thus reflected the variability due to the specific trials assigned to training and testing and, for populations smaller than the full size, the specific units chosen.

#### Fisher Linear Discriminant

The general form of a linear decoding axis is:

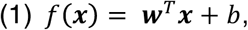

where **w** is an N-dimensional vector (where N is the number of units) containing the linear weights applied to each unit, and b is a scalar value. We fit these parameters using a Fisher Linear Discriminant (FLD), where the vector of linear weights was calculated as:

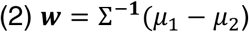

and b was calculated as:

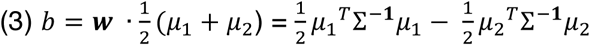

Here *µ*_1_ *and µ*_2_ are the means of the two classes (target matches and distractors, respectively) and the mean covariance matrix is calculated as:

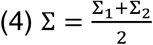

where Σ_1_ and Σ_2_ are the regularized covariance matrices of the two classes. These covariance matrices were computed using a regularized estimate equal to a linear combination of the sample covariance and the identity matrix *I* (Pagan et al., 2016):

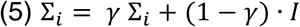

We determined y by exploring a range of values from 0.01 to 0.99, and we selected the value that maximized average performance across all iterations, measured with the cross-validation “regularization” trials set aside for this purpose (see above). We then computed performance for that value of γ with separately measured “test” trials, to ensure a fully cross-validated measure. Because this calculation of the FLD parameters incorporates the off-diagonal terms of the covariance matrix, FLD weights are optimized for both the information conveyed by individual units as well as their pairwise interactions.

To compare FLD performance on correct versus error trials (Fig 6, 7d), we used the same methods described above with the following modifications. First, the analysis was applied to the simultaneously recorded data within each session, and the correlation structure on each trial was kept intact on each resampling iteration. Second, when more than 24 units were available, a subset of 24 units were selected as those with the most task modulation, quantified via the p-value of a one-way ANOVA applied to each unit’s responses (80 conditions * 20 repeats). Finally, on each resampling iteration, each error trial was randomly paired with a correct trial of the same condition and cross-validated performance was performed exclusively for these pairs of correct and error responses. As was the case for the pseudopopulation analysis, training was performed exclusively on correct trials. A mean performance value was computed on each resampling iteration by averaging binary performance outcomes across all possible error trials and their condition-matched correct trial pairs, and averaging across different recording sessions. Mean and standard error of performance was computed as the mean and standard deviation of performance across 100 resampling iterations. Standard error thus reflected error in a manner similar to the pseudopopulation analysis - the variability due to the specific trials assigned to training and testing and, for populations smaller than the full size, the specific units chosen.

In the case of the ranked-FLD (Fig 5c), all units were considered on each resampling iteration, and weights were computed for each unit (with the training data) as described by Equation 2. Weights were then ranked by their magnitude (the absolute values of the signed quantities) and the top N units were selected for different population size N. Finally, both the weights and the threshold were recalculated before cross-validated testing with the training data. In the case of the signed versions of the FLD (which isolated the analysis to target matched enhanced or suppressed units, Fig 7c-d), the process was similar in that all units were considered on each resampling iteration and weights were computed for each unit (with the training data) as described by Equation 2. Weights were then isolated to all of those that were positive “E units” or all that were negative “S units”. Finally, the weights and the threshold were recalculated before cross-validated testing with the training data.

#### Maximum likelihood decoder

As a measure of total IT target match information (combined linear and nonlinear), we implemented the maximum likelihood decoder (Fig 5b) introduced in our previous work (Pagan et al., 2016; Pagan et al., 2013). We began by using the set of training trials to compute the average response r_uc_ of each unit u to each of the 40 conditions c. We then computed the likelihood that a test response k was generated from a particular condition as a Poisson-distributed variable:

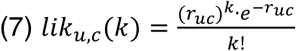

The likelihood that a population response vector was generated in response to each condition was then computed as the product of the likelihoods of the individual units. Next, we computed the likelihood that each test vector arose from the category target match as compared to the category distractor as the product of the likelihoods across the conditions within each category. We assigned the population response to the category with the maximum likelihood, and we computed performance as the fraction of trials in which the classification was correct based on the true labels of the test data.

### Representational similarity

Before computing representational similarity (Fig 9a), the responses of each unit were z-normalized to have a mean of zero and standard deviation of 1. To compute measures of the representational similarity between pairs of population response vectors, the 20 repeated trials for each (of 80) experimental conditions were randomly split into two sets of 10 trials, and the average population response vector was computed. To obtain measures of the noise ceiling, Pearson correlation was computed between many random splits of the data for each of the 80 conditions. The mean across 1000 random splits was computed for each condition and the values were averaged across the splits as well as the 4 target conditions for each image, resulting in 20 correlations values (1 for each image). Fig 9b “Same image and condition” depicts the mean and standard deviation across the 20 images. Measures of the representational similarity between different conditions were computed in a comparable way, by also selecting 10 (of 20) trials before computing the mean population response vectors. To measure the representational similarity between the same objects presented at different transformations, Pearson correlation was computed for all possible pairs of the 5 images corresponding to each object under the conditions of a fixed target. A mean value was computed as the average across 1000 random splits and the pairwise comparison between 1 image and other images containing the same object for each of 20 images, and Fig 9b “Different transforms.” depicts the mean and standard deviation across the 20 images. Fig 9b “Different objects” was computed in a similar manner, but for all possible pairs of one image and the images containing other objects. Finally, Fig 9b “Match versus distractor” was computed in a similar manner, but for all possible pairs of one image presented as a target match (viewing an image while looking for that object as a target) and the three distractor conditions (the three other targets). The same procedures were carried out for Fig 9c and 11b.

### Simulations

To better understand our results, we performed a number of data-based simulations (Fig 11b). Each simulation began by computing the bias-corrected weights for each unit as described above. For the “replication” simulation, we rectified bias-corrected modulations that fell below zero, recomputed the noise-corrected mean spike count responses for each condition, and generated trial variability with an independent Poisson process.

For the “multiplicative, same-sign” simulation, we replaced the target match modulation for each unit with an amount that ensured the population total was distributed proportional to each unit’s total visual modulation (Equation 10), and always reflected as target match enhancement. For the “uniform, mixed-sign” simulation, we replaced each unit’s target match modulation with the same amount, reflected with the sign determined in the original data.

### Statistical tests

When comparing performance between the FLD and maximum likelihood classifier (Fig 5b), we reported *P* values as an evaluation of the probability that differences were due to chance. We calculated *P* values as the fraction of resampling iterations on which the difference was flipped in sign relative to the actual difference between the means of the full data set (for example, if the mean of decoding measure 1 was larger than the mean of decoding measure 2, the fraction of iterations in which the mean of measure 2 was larger than the mean of measure 1).

When evaluating whether each unit had a statistically different response to target matches as compared to distractors (Fig 7a-b, light bars), we recomputed each unit’s modulation index by resampling trials with replacement on n = 1000 resampling iterations. A unit was considered as statistically significant if its resampled modulation indices were flipped in sign from the unit’s actual modulation index less than 0.01% of the resampling iterations. When evaluating whether the single unit modulation indices (Fig 7a-b) were significantly different from zero, we reported P values as computed by a Wilcoxon sign rank test. When evaluating whether the single unit modulation indices computed without repeated distractors (Fig 7b) were significantly different from modulation indices computed with repeated distractors, we reported P values computed via a matched *t* test.

When comparing the representational similarity of different groupings of the IT population response (Fig 7b), we computed a mean Pearson correlation value for each of the 20 images (as described above), and reported P values as the probability that the observed differences in means across the 20 images were due to chance via a two-sample t-test.

### Animal husbandry, enrichment, and care

Monkeys received a nutritionally balanced diet of biscuits as well as daily supplements of fruit and nuts. Monkeys were housed in Allentown cages with space that exceeded the minimums described in the “Guide for Care and Use of Laboratory Animals”. Additionally, monkeys had periodic access to larger playcages that included a variety of enrichment items, such as swings. Monkeys were also provided daily enrichment by social housing when possible and through the introduction of toys, games, and puzzles that involved manipulation to receive food treats. To maintain task motivation, access to water was regulated prior to experimental sessions. Monkeys received a minimum 20 mL/kg of water a day five days a week and a minimum of 40 mL/kg on the other two days. When off study, animals were allowed unrestricted access to water. Animals on regulated access were monitored daily for health status and hydration. Daily hydration status was assessed by body weight, skin turgor, urine and fecal output, and overall demeanor. Following this study, both animals were used in one other neurophysiology study. Following the conclusion of the second study, both animals were euthanized in a manner consistent with the recommendations of the Panel on Euthanasia of the American Veterinary Medical Association, including sedation followed by the introduction of the euthanasia solution Euthasol.

**S1 Dataset**. *IT neural data.* The data include the spike count responses recorded from each monkey, organized into 5-dimensional matrices as units (monkey 1: n = 108; monkey 2: n = 96) x targets (n = 4) x objects (n = 4) x transformations (n = 5) x trials (n = 20). Spikes were counted from 80 to 250 ms, and were extracted from trials with correct responses.

